# Temporal Variation in Mosquito Vector Population Dynamics in Urban Areas

**DOI:** 10.64898/2026.07.13.738234

**Authors:** Shengyang Wang, S. Garrison Carruth, Chalmers Vasquez, John Townsend, Vivek Raman, John-Paul Mutebi, Marco Ajelli, André B.B. Wilke

## Abstract

Locally acquired arboviral infections are an increasing public health concern in the United States. *Aedes aegypti* and *Culex quinquefasciatus*, vectors of dengue virus and West Nile virus, respectively, are established in regions where local transmission has been reported. Seasonal variation in mosquito abundance affects vector density, human-vector contact, and arbovirus transmission risk. The aim of this study is to investigate seasonal variations in *Ae. aegypti* and *Cx. quinquefasciatus* population dynamics in Miami-Dade County, Florida; Maricopa County, Arizona; and Clark County, Nevada. Monthly relative abundance, average mosquitoes collected per trap-night, normalized abundance ratios, and seasonal-trend decomposition were used to evaluate species- and location-specific seasonal patterns. Mosquito population dynamics differed by species and location. In Miami-Dade County, the two species showed seasonal turnover, with *Cx. quinquefasciatus* peaking during winter and spring and *Ae. aegypti* peaking during summer. In Maricopa and Clark counties, both species peaked primarily between August and September. Species dominance also differed by site, with *Ae. aegypti* more abundant than *Cx. quinquefasciatus* in Maricopa County and *Cx. quinquefasciatus* more abundant than *Ae. aegypti* in Clark and Miami-Dade counties. Seasonal decomposition showed that *Ae. aegypti* peaked earlier in Miami-Dade County than in Maricopa and Clark counties, whereas *Cx. quinquefasciatus* showed a spring peak in Miami-Dade County and bimodal seasonal patterns in Maricopa and Clark counties. These results suggest that mosquito population dynamics are species- and location-specific and support the use of local surveillance data to guide the timing of mosquito control and arbovirus preparedness.

## Introduction

Locally acquired arboviral infections are an increasing public health concern in the United States. West Nile virus (WNV) is endemic across much of the continental United States and represents the most widespread mosquito-borne viral disease in the country (Centers for Disease Control and Prevention 2026). Nonetheless, its burden is substantially underestimated, with one study estimating approximately 7 million WNV infections in the United States from 1999 to 2016 (Ronca et al. 2019), compared with 61,013 reported cases from 1999 to 2024 (Centers for Disease Control and Prevention 2025). In contrast, locally acquired dengue virus (DENV) infections remain less common in the continental United States but have been increasingly reported in recent years. From 2010 to 2024, locally acquired DENV infections were reported in California, Arizona, Texas, North Carolina, West Virginia, New York, and Florida (Centers for Disease Control and Prevention 2024). Although reported locally acquired dengue cases remain relatively low in the continental United States, the true burden is underestimated, with one CDC study estimating a reporting rate of 1.0%–4.8% (Shankar et al. 2018). WNV and DENV are transmitted by mosquito vectors, including *Culex quinquefasciatus* and *Aedes aegypti*, respectively. Both species are established and expanding their range across the United States, including parts of California, Arizona, and Nevada (Monaghan et al. 2019; Gorris et al. 2021).

Temporal variability in mosquito population dynamics drives both vector distribution and density (Lana et al. 2014; Manore et al. 2017; Murdock et al. 2017; Andreo et al. 2021). These dynamics frequently follow seasonal cycles, with periods of elevated abundance and expanded range increasing human-vector contact rates and, consequently, amplifying the risk of arbovirus transmission (Espinosa et al. 2016; Dzul-Manzanilla et al. 2021; Zahid et al. 2023; Wilke et al. 2023). Characterizing population dynamics, therefore, supports the implementation of mosquito control strategies and outbreak preparedness by informing the timing of preventive interventions ahead of seasonal increases in abundance and range (Lizzi et al. 2014; Ventura et al. 2024; Harp et al. 2025). However, the extent to which the population dynamics of vector mosquito species differ among urban areas across the United States remains unclear. To address this gap, the aim of this study is to compare the seasonal patterns of *Ae. aegypti* and *Cx. quinquefasciatus* across three urban areas with distinct climatic and socioecological contexts: Miami-Dade County, Florida; Maricopa County, Arizona; and Clark County, Nevada.

## Methods

### Study sites

In this study, we selected Miami-Dade County, Florida; Maricopa County, Arizona; and Clark County, Nevada, due to their distinct climatic and socioecological profiles, as well as recent reports of locally acquired DENV and WNV transmission (Centers for Disease Control and Prevention 2025) Miami-Dade County has a tropical climate, with consistently warm temperatures and high humidity. In contrast, Maricopa County is characterized by a hot desert climate, with intense summer heat, persistent drought, and limited water resources. However, Maricopa undergoes yearly increases in precipitation associated with the North American Monsoon, which increases regional humidity during summer months (National Oceanic and Atmospheric Administration 2021). Clark County, located in the Mojave Desert, exhibits a subtropical hot desert climate with extremely high summer temperatures, lower annual precipitation, sparse vegetation, and pronounced diurnal temperature variation (Kottek et al. 2006).

### Data Collection

Mosquito collections were conducted through existing surveillance systems across the study sites. Mosquito surveillance systems had similar traps, deployment frequency, and spatial resolution. We analyzed specimens collected from two trap types: BG-Sentinel and CDC light traps baited with CO_2_. Within each study site, traps were deployed for 24 hours per sampling event (i.e., one trap-night). Collected mosquitoes were morphologically identified to species level using the standard taxonomic keys (Darsie, Jr. and Morris 2000). A descriptive analysis of the relative abundance of *Ae. aegypti* and *Cx. quinquefasciatus* in Miami-Dade County has been reported previously (Vasquez et al. 2026), while the present study aims at a multi-site comparison, which includes a signal decomposition analysis.

Host-seeking *Ae. aegypti* and *Cx. quinquefasciatus* female mosquitoes were collected weekly from January 2020 to December 2024 in Miami-Dade and Maricopa Counties. In Clark County, collections were conducted from April to October from 2020 to 2024 due to cold winter temperatures that substantially limit mosquito activity in desert environments during the colder months (November to March). Although a small subset of traps collected male mosquitoes, they were considered accidental catches and excluded from the analysis.

### Data analysis

For each study site, we recorded the relative abundance of *Ae. aegypti* and *Cx. quinquefasciatus.* To ensure sampling consistency across sites, traps deployed fewer than 30 times per year (<30 trap-nights) or more than 54 times per year (>54 trap-nights) were removed from the analyses. These thresholds were selected to retain trap locations sampled approximately weekly, while excluding locations sampled inconsistently or more frequently than the standard surveillance schedule.

Then, we calculated the monthly trap-nights (i.e., the total number of traps deployed for 24 hours per month) to quantify sampling effort, as well as the average number of *Ae. aegypti* and *Cx. quinquefasciatus* collected per trap-night per month. We further normalized the data by calculating a ratio of the monthly relative abundance divided by the maximum monthly relative abundance recorded during the study period, allowing for the comparison of mosquito seasonal dynamics across study sites. All the data were analyzed using R (version 4.5.1).

### Signal Decomposition Analysis

To disentangle long-term trends from recurring seasonal variation and residual variation in mosquito relative abundance, we applied Seasonal-Trend decomposition based on Loess (STL) (Cleveland RB 1990). For each species and study site, the time series was defined as the monthly average number of mosquitoes collected per trap-night, thus accounting for variation in sampling effort. STL employs a locally weighted regression to decompose data into three components:

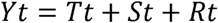

where *Y_t_* is the monthly number of mosquitoes collected per trap-night at month *t*, *T_t_* is the trend component that reflects long-term population dynamics, *S_t_* is the seasonal component, and *R_t_* is the residual component. STL decomposition was performed using the *stl*()function in R (version 4.5.1), with a periodic seasonal window of 12 months to represent the recurring seasonal structure. For Clark County, where surveillance was conducted from April to October, months without collections were retained as zeros to preserve a 12-month annual temporal structure.

## Results

### Relative abundance: within-site species comparisons

The relative abundance of *Ae. aegypti* and *Cx. quinquefasciatus* differed by species and study site, with consistent differences in both total monthly collections and trap-night-adjusted abundance (Fig. 1 and 2).

**Figure 1.**
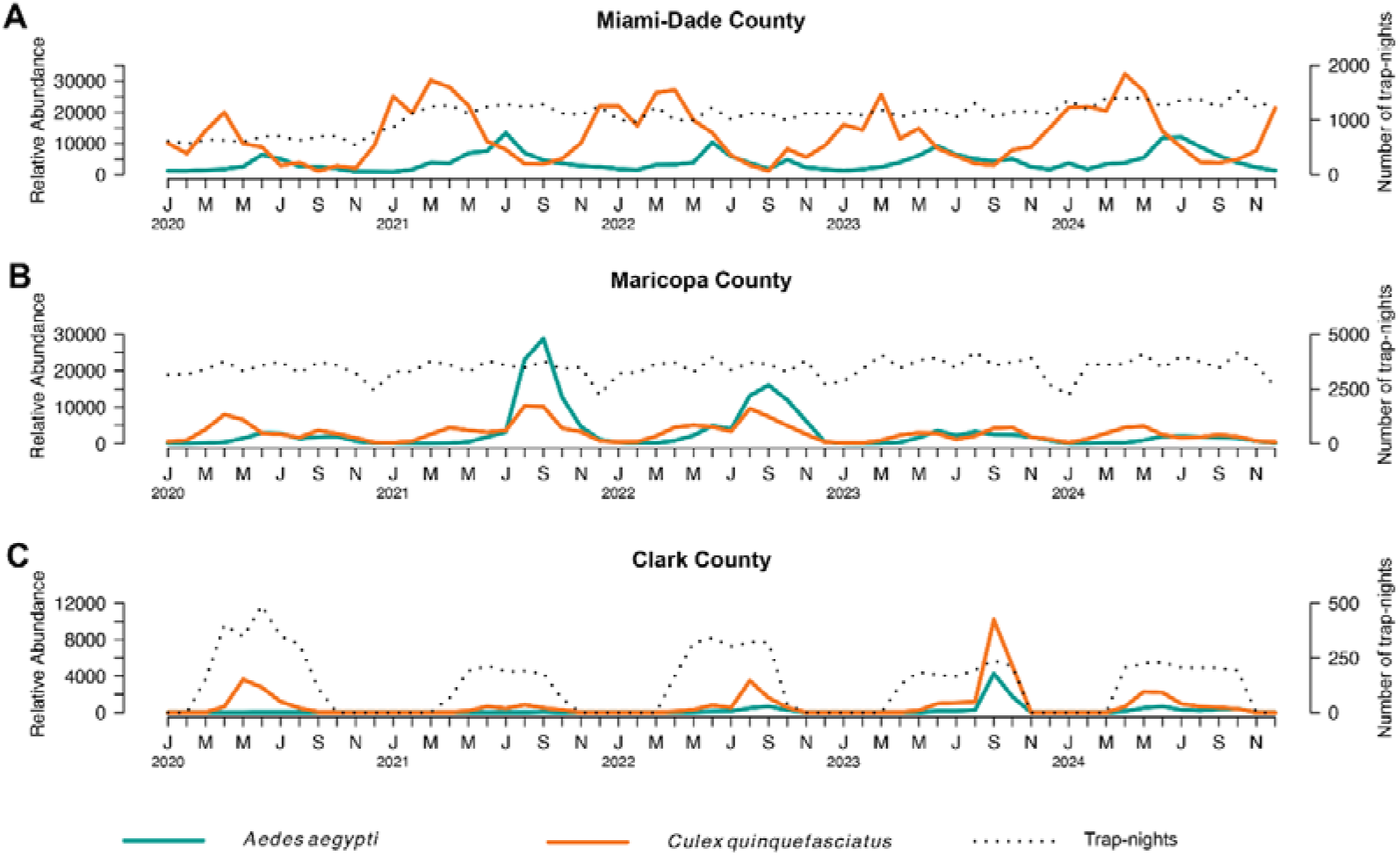
**(A)** Number of collected *Ae. aegypti* and *Cx. quinquefasciatus* per month from 2020 to 2024 in Miami-Dade County, Florida. The dotted line shows the monthly number of trap-nights (scale on the right axis). **(B)** Same as A, but for Maricopa County, Arizona. **(C)** Same as A, but for Clark County, Nevada.

**Figure 2.**
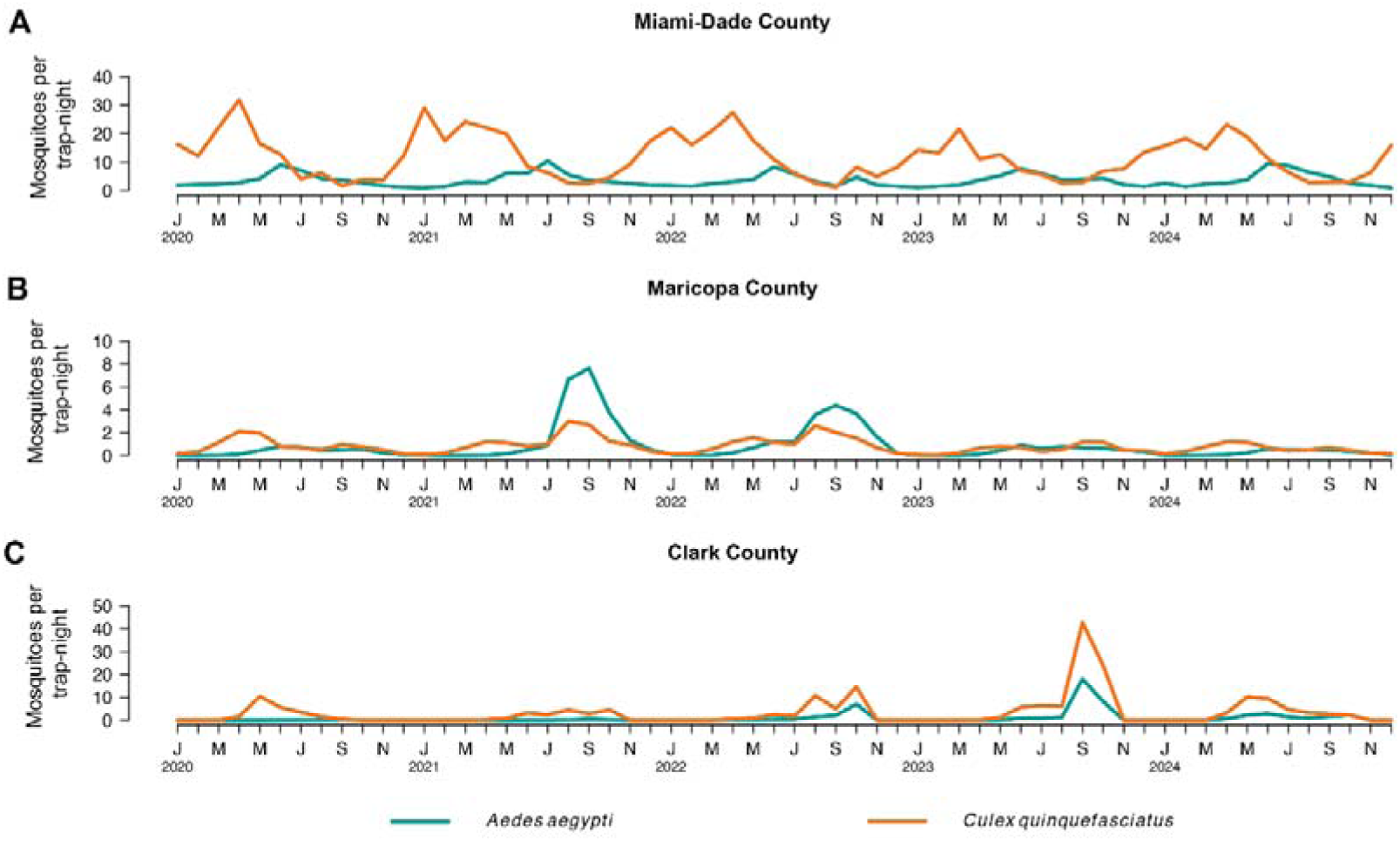
**(A)** Number of collected *Ae. aegypti* and *Cx. quinquefasciatus* per month per trap night from 2020 to 2024 in Miami-Dade County, Florida. **(B)** Same as A, but for Maricopa County, Arizona. **(C)** Same as A, but for Clark County, Nevada.

In Miami-Dade County, *Cx. quinquefasciatus* was consistently more abundant than *Ae. aegypti* throughout the study period. Both species showed seasonal patterns, but peak abundance differed by species. *Aedes aegypti* relative abundance increased from June through August (reaching a maximum of 13,356 specimens in July 2021) and declined to its lowest levels between December and January. In contrast, *Cx. quinquefasciatus* relative abundance increased from December through April (reaching a maximum of 32,213 specimens in April 2024) and declined to its lowest levels in September. Sampling effort, measured as the number of trap-nights per month, increased from approximately 650 trap-nights in 2020 to 1,200 trap-nights in 2021 and remained relatively stable thereafter (Fig. 1A). After accounting for sampling effort, yearly peak relative abundance ranged from 7.70 to 10.42 mosquitoes per trap-night for *Ae. aegypti* and from 21.59 to 31.80 mosquitoes per trap-night for *Cx. quinquefasciatus* (Fig. 2A).

In Maricopa County, *Ae. aegypti* was consistently more abundant than *Cx. quinquefasciatus*. Both species showed seasonal patterns, although these patterns were less distinct than those observed in Miami-Dade County. *Aedes aegypti* relative abundance peaked in September 2021 and September 2022, with 28,814 and 16,041 specimens collected, respectively, and declined to its lowest levels between December and January. *Culex quinquefasciatus* followed a similar seasonal pattern, peaking in August 2021 and August 2022, with 10,342 and 9,621 specimens collected, respectively, and declining to its lowest levels between December and January. Sampling effort remained stable throughout the study period, with an average of 3,468 trap-nights per month (Fig. 1B). After accounting for sampling effort, *Ae. aegypti* peaks reached 7.64 and 4.40 mosquitoes per trap-night in September 2021 and September 2022, respectively, whereas *Cx. quinquefasciatus* remained below three mosquitoes per trap-night throughout the study period, reaching 2.98 and 2.60 mosquitoes per trap-night in August 2021 and August 2022, respectively (Fig. 2B).

In Clark County, *Cx. quinquefasciatus* was consistently more abundant than *Ae. aegypti*. Both species showed similar seasonal patterns, with peak abundance occurring between August and September. Relative abundance was low during 2020 and 2021 for both species and increased during 2022 and 2023. *Aedes aegypti* increased from 657 specimens collected in 2022 to 4,279 specimens in 2023. *Culex quinquefasciatus* also increased during this period, reaching 10,158 specimens collected in 2023. Sampling effort remained relatively stable from April through September, when the surveillance system was active, with an average of 224 trap-nights per month (Fig. 1C). After accounting for sampling effort, trap-night-adjusted abundance was highest in September 2023, when *Cx. quinquefasciatus* reached 42.68 mosquitoes per trap-night and *Ae. aegypti* reached 17.98 mosquitoes per trap-night; in 2024, peak abundance declined to 10.22 and 2.82 mosquitoes per trap-night, respectively (Fig. 2C).

### Seasonal patterns: within-site species comparisons

The normalized abundance ratios and STL-derived seasonal components showed distinct species-specific seasonal structures across the three study sites (Fig. 3). In Miami-Dade County, the two species showed the clearest seasonal offset, with *Cx. quinquefasciatus* characterized by spring-dominated activity and *Ae. aegypti* by summer-dominated activity, resulting in seasonal turnover between the two species (Fig. 3A,B). In Maricopa County, seasonal activity was partially synchronized in late summer, but the STL decomposition distinguished a unimodal September peak for *Ae. aegypti* from a bimodal pattern for *Cx. quinquefasciatus*, with elevated activity in spring and again in August–September (Fig. 3C,D). In Clark County, the seasonal structure was less consistent across years, with a weakly bimodal pattern and more episodic peaks for both species, although the strongest seasonal activity occurred in September–October (Fig. 3E,F).

**Figure 3.**
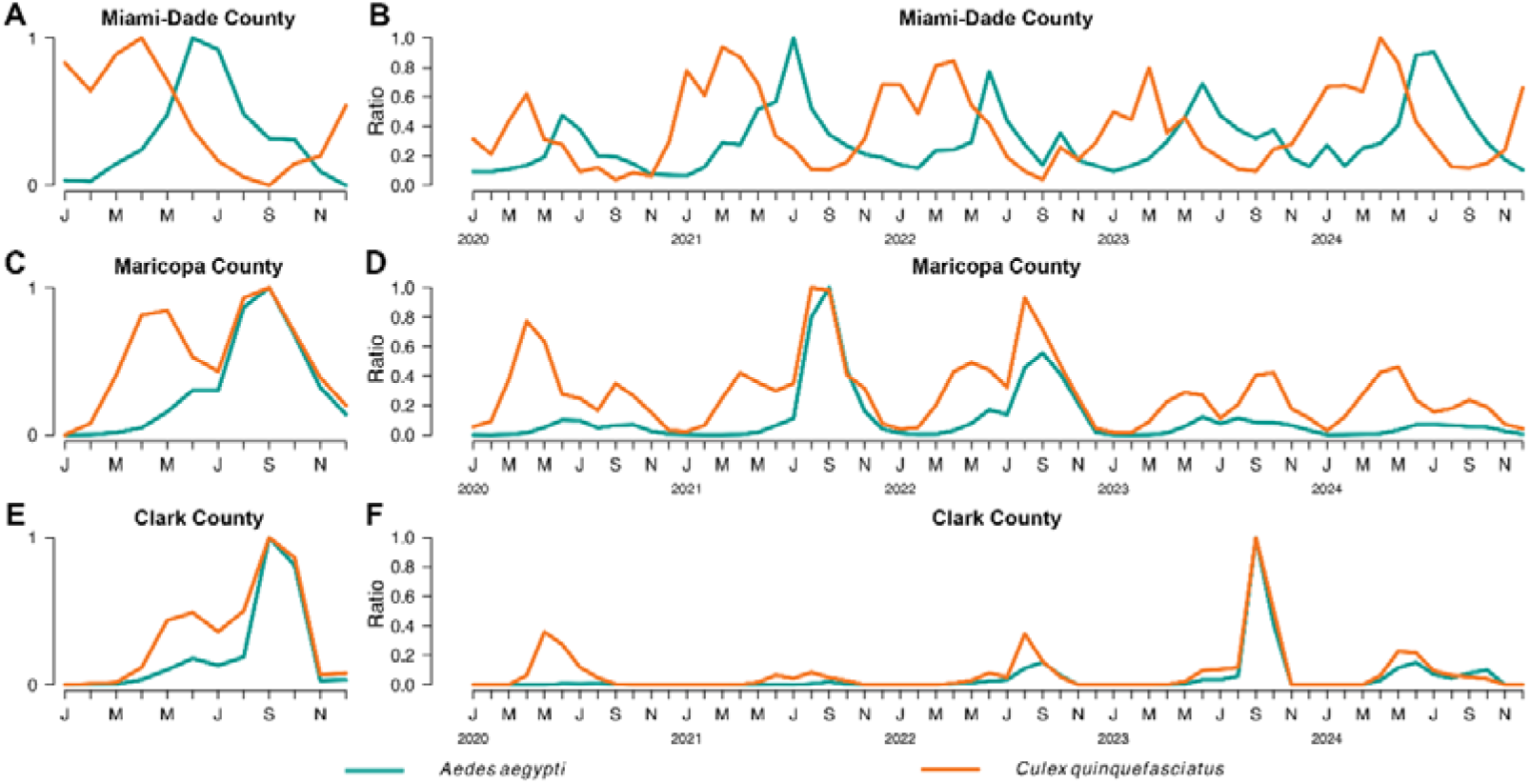
**(A)** STL-derived seasonal components for *Ae. aegypti* and *Cx. quinquefasciatus* from 2020 to 2024 in Miami-Dade County, Florida. **(B)** Monthly normalized abundance ratios for *Ae. aegypti* and *Cx. quinquefasciatus* from 2020 to 2024 in Miami-Dade County, Florida. **(C)** Same as A, but for Maricopa County, Arizona. **(D)** Same as B, but for Maricopa County, Arizona. **(E)** Same as A, but for Clark County, Nevada. **(F)** Same as B, but for Clark County, Nevada.

### Relative abundance: across-site comparisons within species

Relative abundance differed across study sites for both *Ae. aegypti* and *Cx. quinquefasciatus*, whether expressed as total monthly collections or as collections per trap-night (Figs. 4 and 5, respectively).

**Figure 4.**
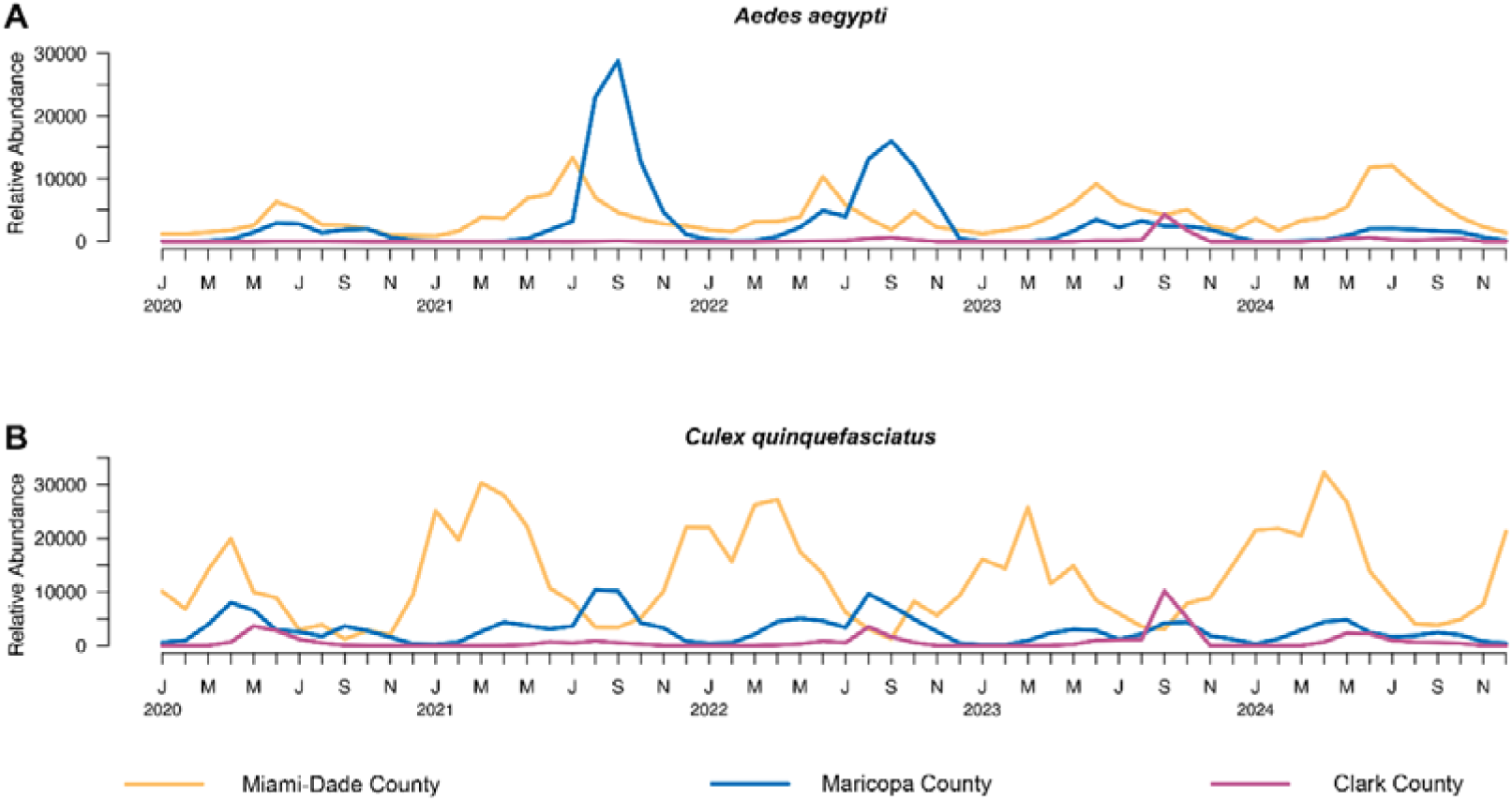
**(A)** Monthly relative abundance of *Ae. aegypti* from 2020 to 2024 in Miami-Dade County, Florida; Maricopa County, Arizona; and Clark County, Nevada. **(B)** Same as A, but for *Cx. quinquefasciatus*.

**Figure 5.**
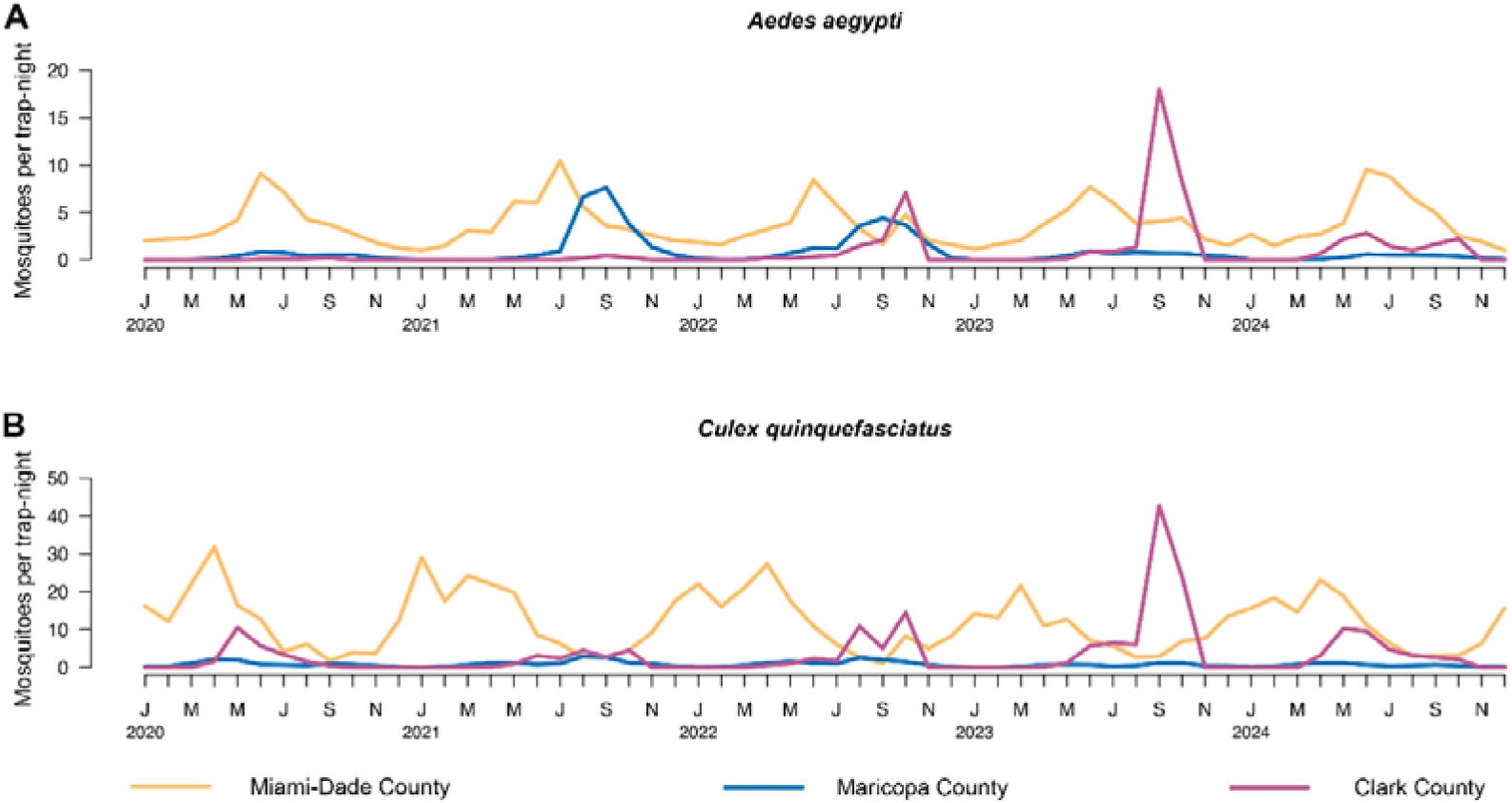
**(A)** Number of collected *Ae. aegypti* per month per trap-night from 2020 to 2024 in Miami-Dade County, Florida; Maricopa County, Arizona; and Clark County, Nevada. **(B)** Same as A, but for *Cx. quinquefasciatus*

For *Ae. aegypti*, Maricopa County showed the highest relative abundance, with peaks of 28,814 specimens in September 2021 and 16,041 specimens in September 2022. Miami-Dade County had lower relative abundance than Maricopa County but showed more consistent seasonal recurrence, with monthly values ranging from 869 to 13,356 specimens. Clark County had the lowest overall relative abundance, although abundance increased over time and reached 4,279 specimens in September 2023. After accounting for sampling effort, Miami-Dade County generally had higher relative abundance per trap night than Maricopa County with more consistent peak values in different years. In Clark County, trap-night-adjusted abundance remained low during 2020 and 2021 but reached 17.98 mosquitoes per trap-night in September 2023, the highest value observed for *Ae. aegypti* across the three study sites (Figs. 4A and 5A).

For *Cx. quinquefasciatus*, Miami-Dade County showed the highest relative abundance, with monthly values ranging from 1,227 to 32,213 specimens. Maricopa County had lower relative abundance, with peaks of 10,342 and 9,621 specimens in August 2021 and August 2022, respectively. Clark County had the lowest overall relative abundance but increased over time, reaching 10,158 specimens in September 2023 before declining in 2024. After accounting for sampling effort, *Cx. quinquefasciatus* relative abundance was much higher in Miami-Dade County than in the other two study sites, with peak values consistently over 20 collected mosquitoes per trap-night in Miami-Dade. Clark County showed a high variability between seasons, while consistently lower than 3 collected mosquitoes per trap night were found in Maricopa County (Figs. 4B and 5B).

### Seasonal patterns: across-site comparisons within species

The normalized abundance ratios and STL-derived seasonal components showed that seasonal timing differed across study sites within each species (Fig. 6). For *Ae. aegypti*, Miami-Dade County showed earlier seasonal activity, with a main peak in June, whereas Maricopa and Clark counties showed later activity, with peaks concentrated in September, particularly during years with higher abundance (Fig. 6A,B). For *Cx. quinquefasciatus*, Miami-Dade County showed a unimodal spring pattern, peaking in April, whereas Maricopa and Clark counties showed bimodal seasonal structure, with elevated activity during spring or early summer and again in late summer (Fig. 6C,D). This bimodal structure was more consistent in Maricopa County and less stable across years in Clark County.

**Figure 6.**
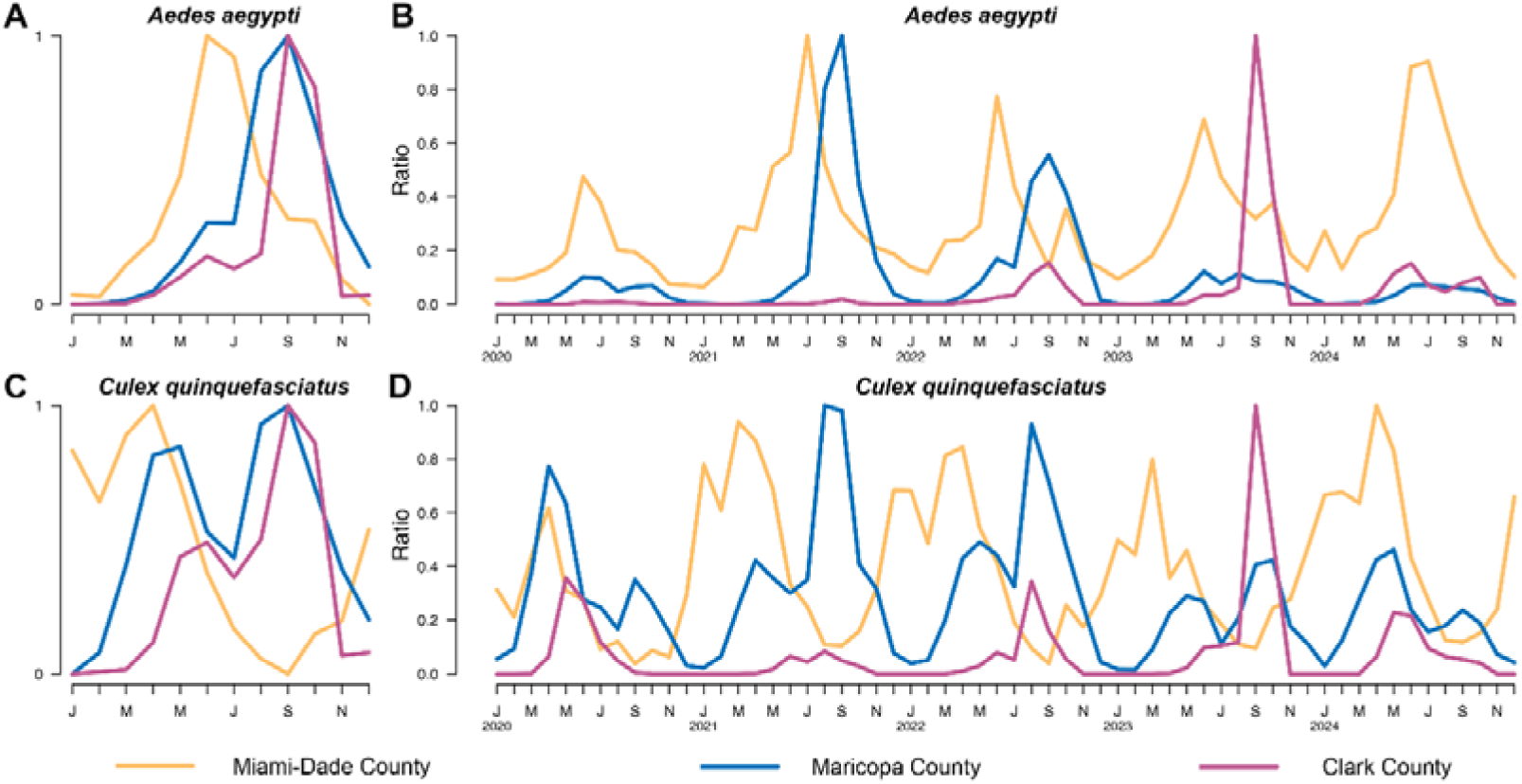
**(A)** STL-derived seasonal components for *Ae. aegypti* from 2020 to 2024 in Miami-Dade County, Florida; Maricopa County, Arizona; and Clark County, Nevada. **(B)** Monthly normalized abundance ratios for *Ae. aegypti* from 2020 to 2024 in Miami-Dade County, Florida; Maricopa County, Arizona; and Clark County, Nevada. **(C)** Same as A, but for *Cx. quinquefasciatus*. **(D)** Same as B, but for *Cx. quinquefasciatus*.

## Discussion

Arboviral infections remain a public health concern globally and in the United States. Understanding the temporal population dynamics of mosquito vectors is important for anticipating periods of elevated vector abundance and informing mosquito control strategies. In this study, we found location-dependent heterogeneity in mosquito population dynamics across three urban areas, with distinct within- and between-species patterns. These findings support the use of local surveillance data to guide the timing of mosquito surveillance and control.

In Miami-Dade County, *Ae. aegypti* and *Cx. quinquefasciatus* showed a well-defined seasonal turnover. *Cx. quinquefasciatus* was more abundant earlier in the year, whereas *Ae. aegypti* increased later and peaked during summer months. The consistent offset between the seasonal peaks of *Cx. quinquefasciatus* and *Ae. aegypti* suggests that mosquito population dynamics in Miami-Dade County were relatively stable during the study period. These patterns are relevant for arbovirus preparedness because they indicate that periods of elevated vector abundance differ between DENV and WNV vectors within the same geographic area.

A different pattern was observed in Maricopa and Clark counties. After accounting for sampling effort, the average number of mosquitoes collected per trap-night was lower in Maricopa and Clark counties than in Miami-Dade County for both species. In Maricopa County, *Ae. aegypti* was more abundant than *Cx. quinquefasciatus*, and both species reached their highest relative abundance in late summer, primarily in September. In Clark County, both species showed low activity during 2020 and 2021, followed by increases in 2022 and a marked peak in 2023. The September 2023 peak in Clark County yielded the highest trap-night-adjusted abundance observed for both *Ae. aegypti* and *Cx. quinquefasciatus* across all study sites, which may have been influenced by the environmental conditions associated with Tropical Storm Hilary (National Oceanic and Atmospheric Administration 2023). In 2024, the average number of mosquitoes collected per trap-night declined. These results indicate that mosquito populations in desert urban environments may remain low for extended periods and increase during specific seasonal windows when conditions become conducive.

The signal decomposition analysis further showed that seasonal structure differed by species and location. *Aedes aegypti* peaked earlier in Miami-Dade County, with maximum seasonal activity around June, whereas *Ae. aegypti* populations in Maricopa and Clark counties peaked later, primarily in September. In contrast, *Cx. quinquefasciatus* showed a unimodal spring pattern in Miami-Dade County and bimodal seasonal patterns in Maricopa and Clark counties, with activity occurring during spring or early summer and again in late summer. These differences suggest that the same vector species can follow different seasonal trajectories across urban environments. Regional climate and local habitat conditions may contribute to differences in the timing and duration of vector activity. Previous studies have shown that mosquito population dynamics and seasonality are influenced by climatic factors, including temperature and rainfall (Marini et al. 2016; Beck-Johnson et al. 2017; Whittaker et al. 2023; Blanco-Sierra et al. 2024). Future studies could evaluate how these environmental variables contribute to mosquito population dynamics in Miami-Dade, Maricopa, and Clark counties.

Consistent with previous research (Biteye et al. 2018; Whittaker et al. 2022; Rakotoarison et al. 2025), we found seasonal and geographic heterogeneity in mosquito population dynamics. These site-specific differences support the use of local seasonal surveillance thresholds rather than uniform control schedules across regions. They also suggest that arbovirus preparedness should consider the timing of vector abundance separately for each vector species. In Miami-Dade County, interventions targeting *Cx. quinquefasciatus* may need to begin before winter, whereas interventions targeting *Ae. aegypti* should be intensified before summer. In Maricopa and Clark counties, *Ae. aegypti* control may be most relevant before late-summer increases, whereas *Cx. quinquefasciatus* control may require attention during both spring and late summer because of its bimodal seasonal pattern.

We relied on existing mosquito surveillance systems for cross-site comparison. Although trap deployment frequency and spatial resolution were similar across sites, differences in trap placement and mosquito habitat availability may have influenced collections. The analysis focused on adult female abundance and did not include infection rates, vector competence, human behavior, or host availability, all of which influence arbovirus transmission risk. Environmental variables such as temperature and rainfall were also not included, limiting inference about the drivers of the observed patterns. Despite these limitations, identifying species-specific and location-specific heterogeneity in mosquito population dynamics can inform mosquito surveillance and control systems by supporting the timing of public health actions and vector control interventions (Tsai et al. 2012; Fotakis et al. 2022; Baafi and Hurford 2025).

## Conclusion

This study provides evidence that *Ae. aegypti* and *Cx. quinquefasciatus* population dynamics vary across urban areas. The timing, magnitude, and seasonal structure of vector abundance differed by geographic location and species. These results support the need for localized mosquito surveillance and control strategies. Future work could integrate mosquito abundance with environmental covariates, spatial habitat data, arbovirus testing, and human case data to identify the ecological drivers of seasonal vector dynamics and improve outbreak preparedness.

## Data Availability

Data from Miami-Dade, Maricopa, and Clark counties analyzed in this study were obtained from publicly supported surveillance programs. Although such data are public property and may ultimately become part of the public record, access is not automatically unrestricted or immediate. Individuals seeking access to the surveillance data used in this study must obtain permission from the appropriate public health authority to ensure compliance with applicable public-records statutes, data-use policies, and privacy protections. Researchers interested in obtaining the dataset may submit a formal Public Records Request to: Miami-Dade County Mosquito Control Division, available at: https://www.miamidade.gov/global/solidwaste/mosquito/contact-mosquito-control.page.

Maricopa County Environmental Services, Department of Vector Control Division, available at: https://publicrecordsrequest.maricopa.gov/nonCommercial?d=133

Southern Nevada Health District, Environmental Health, Public Accommodations, and Mosquito Disease Surveillance. available at: https://www.southernnevadahealthdistrict.org/news-info/records-request/public-records-request/

